# Temporal learning among prefrontal and striatal ensembles

**DOI:** 10.1101/2020.02.28.970053

**Authors:** Eric Emmons, Gabriela Tunes-Chiuffa, Jeeyu Choi, R. Austin Bruce, Matthew A. Weber, Youngcho Kim, Nandakumar S. Narayanan

## Abstract

Behavioral flexibility requires the prefrontal cortex and striatum. Here, we investigate neuronal ensembles in the medial frontal cortex (MFC) and the dorsomedial striatum (DMS) during one form of behavioral flexibility: learning a new temporal interval. We studied corticostriatal neuronal activity as rodents trained to respond after a 12-second fixed interval (FI12) learned to respond at a shorter 3-second fixed interval (FI3). On FI12 trials, we discovered time-related ramping was reduced in the MFC but not in the DMS in two-interval vs. one-interval sessions. We also found that more DMS neurons than MFC neurons exhibited differential interval-related activity on the first day of two-interval performance. Finally, MFC and DMS ramping was similar with successive days of two-interval performance but DMS temporal decoding increased on FI3 trials. These data suggest that the MFC and DMS play distinct roles during temporal learning and provide insight into corticostriatal circuits.

## Introduction

Behavioral flexibility requires learning to adapt to uncertainty. Two forebrain structures critical for flexibility are the prefrontal cortex and striatum (Fuster, 2008; Kehagia et al., 2010). Prefrontal cortical neurons densely innervate the striatum (Gabbott et al., 2005; Wall et al., 2013) and disruptions of either structure profoundly impact the learning of new goals, rules, and strategies (Hart et al., 2018; Ragozzino, 2007). Dysfunctional corticostriatal circuits and connectivity are implicated in a range of psychiatric and neurological disorders (Deutch, 1993; Shepherd, 2013). However, the relative roles of prefrontal and striatal networks during behavioral flexibility are unclear.

One task that provides an ideal window into behavioral flexibility is interval timing, which requires participants to estimate an interval of several seconds via a motor response. Across species, interval timing requires the prefrontal cortex and striatum (Coull et al., 2011; Dallérac et al., 2017; Emmons et al., 2017, 2016; Matell and Meck, 2004; Merchant and de Lafuente, 2014). Work from our group and others has shown that both prefrontal and striatal neurons encode temporal information via ‘time-related ramping’ activity—or monotonic changes in firing rate over a temporal interval (Bakhurin et al., 2017; Donnelly et al., 2015; Emmons et al., 2017; Kim et al., 2018; Narayanan, 2016; Wang et al., 2018). Our past work suggested that ramping activity in neurons of the medial frontal cortex (MFC) and the dorsomedial striatum (DMS) is very similar, with ∼40% of neurons in each area exhibiting such activity (Emmons et al., 2017). We have also found that MFC inactivation attenuates DMS ramping (Emmons et al., 2019, 2017) and that MFC stimulation is sufficient to increase DMS ramping (Emmons et al., 2019). These data suggest that DMS ramping is closely linked to MFC ramping and suggest the hypothesis that MFC and DMS ensembles respond similarly as animals learn new temporal intervals. By contrast, recordings from primate lateral prefrontal cortex and caudate indicate that striatal ensembles encode stimulus-response associations earlier than prefrontal ensembles, leading to the hypothesis that prefrontal and striatal ensembles play differential roles during learning (Antzoulatos and Miller, 2011; Histed et al., 2009; Pasupathy and Miller, 2005).

We tested these hypotheses by recording MFC and DMS activity in rodents as they learned to respond to a new interval. Specifically, rodents previously trained to perform a 12-second fixed-interval task learned a new version of the task that included two fixed-intervals—3 seconds and 12 seconds. We report three main results. First, time-related ramping activity in the MFC decreased on 12-second interval trials during two-interval sessions compared to one-interval sessions, whereas activity in the DMS on 12-second interval trials did not change. Second, DMS neurons were more likely to have distinct firing patterns during the 12-second vs. 3-seconds interval than those of the MFC, but this interval-related activity normalized between MFC and DMS over subsequent days of two-interval performance. Finally, MFC and DMS ramping did not change consistently over subsequent two-interval sessions, but temporal decoding by DMS ensembles improved for FI3 trials. These data suggest that the MFC and DMS play distinct roles during temporal learning.

## Methods

### Rodents

All procedures were approved by the University of Iowa IACUC, and all methods were performed in accordance with the relevant guidelines and regulations (protocol #7072039). Seven male Long-Evans rats were trained on the 12-second fixed-interval timing task (FI12) according to procedures described in detail previously (Emmons et al., 2017, 2016). In brief, the rats were autoshaped to press a lever for water reward using a fixed-ratio task before being trained on 12-second fixed-interval timing (FI12). Trials began with the presentation of a house light, and the first response made after 12 seconds resulted in the delivery of a water reward, a concurrent click, and termination of the house light (Fig. 1A; video S1). Responses made before the interval ended were unreinforced. Trials were separated by a randomly chosen 6-, 8-, 10-, or 12-second intertrial interval. After animals behaved consistently, the MFC and DMS were each implanted with recording electrodes (Fig. 1B; see below). Animals were then acclimatized to the recording procedures and recordings were made during behavior in the FI12 task (Day 0). The following day, an additional 3-s interval (FI3) was added to the task and cued by a light distinct from the one used to indicate FI12. Behavior and simultaneous neuronal activity in the MFC and DMS were recorded over the following three days (Day 1, Day 2, and Day 3). Some data from subsequent recording sessions in these rodents were included in prior manuscripts (Emmons et al., 2017, 2016).

**Figure 1:**
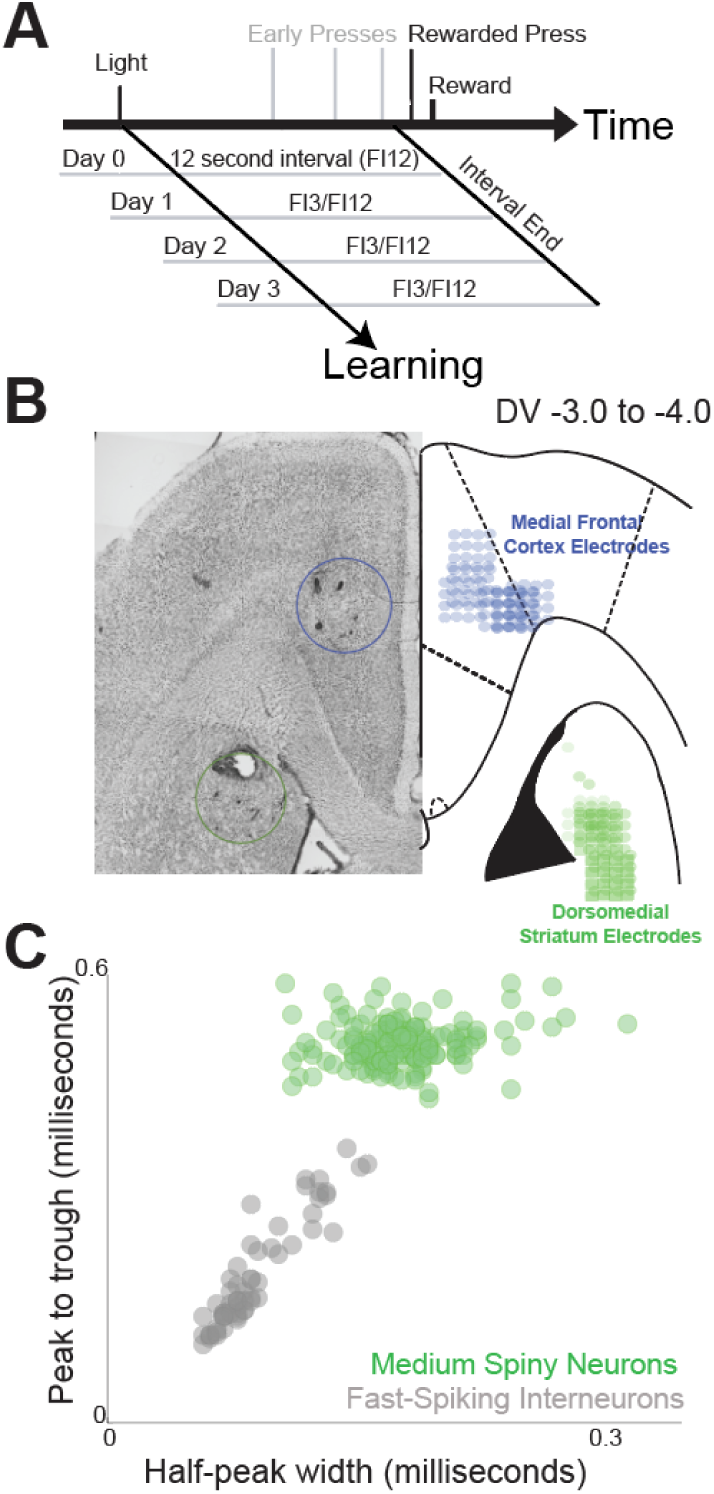
Fixed-interval timing tasks and recording locations. A) On Day 0, rodents performed fixed-interval timing tasks in which a reward was given for the first lever press after a 12-second interval (FI12). Interval start was cued by a house light, motivation was a liquid reward, and presses before interval end were unreinforced. On Day 1, a second, shorter 3-second interval (FI3) was introduced and cued by a distinct light. FI3 trials were randomly intermixed with FI12 trials. Recordings were performed for two days following the initial two-interval performance (Day 2 and Day 3). B) Animals were implanted with neuronal ensemble arrays targeting the medial frontal cortex (MFC) and dorsomedial striatum (DMS). C) MSNs within the DMS were identified based on waveform shape.

### Surgical and histological procedures

Animals were anesthetized using ketamine (100 mg/kg) and xylazine (10 mg/kg), and a surgical level of anesthesia was maintained using ketamine supplements (10 mg/kg). Craniotomies were drilled above the left MFC and left DMS and four holes were drilled for skull screws, which were connected to electrode recording arrays via a separate ground wire. Microelectrode arrays were composed of 4×4 50-μm stainless steel wires (250 μm between wires and rows; impedance measured *in vitro* at ∼400 kΩ; Plexon, Dallas, TX). These arrays were positioned in the MFC (coordinates from bregma: AP +3.2, ML ±1.2, DV −3.6 @ 12° in the lateral plane) and the DMS (coordinates from bregma: AP +0.0, ML ±4.2, DV −3.6 @ 12° in the posterior or lateral plane) while recording neuronal activity to verify that implantation was in correct brain area. The craniotomy was sealed with cyanoacrylate (‘SloZap’, Pacer Technologies, Rancho Cucamonga, CA), and the reaction was accelerated by ‘ZipKicker’ (Pacer Technologies) and methyl methacrylate (AM Systems, Port Angeles, WA). Rats recovered for one week before being acclimatized to behavioral and recording procedures.

Following these experiments, the rats were anesthetized and sacrificed by injection with 100 mg/kg sodium pentobarbital and transcardially perfused with 4% formalin. Brains were post-fixed in a solution of 4% formalin and 20% sucrose before being sectioned on a freezing microtome. Brain slices were mounted on Superfrost Plus microscope slides and stained for cell bodies using either DAPI or Cresyl violet. Histological reconstruction was completed using postmortem analysis of electrode placement by slide-scanning fluorescent microscopy (Olympus).

### Neurophysiological recordings and neuronal analyses

Neuronal ensemble recordings were made using a multi-electrode recording system (Plexon). In each animal, one electrode without single units was reserved for local referencing, yielding 15 electrodes per animal. Offline Sorter (Plexon) was used to analyze the signals after the experiments and to remove artifacts. Spike activity was analyzed for all cells that fired at rates above 0.1 Hz. Principal component analysis (PCA) and waveform shape were used for spike sorting. DMS neurons were classified as either medium spiny neurons (MSNs) or interneurons based on peak-to-trough ratio and the spike half-peak width of spike waveforms (Fig. 1C; Berke, 2011). Single units were defined as those 1) having a consistent waveform shape, 2) being a separable cluster in PCA space, and 3) having a consistent refractory period of at least 2 ms in interspike interval histograms.

### Statistics

Basic analyses were performed via ANOVA, Wilcoxon rank-sum tests, and Cohen’s D. As in our prior work, we quantified temporal control of action during fixed-interval performance in two ways. We calculated the curvature of time-response histograms (Emmons et al., 2019; Fry et al., 1960; Narayanan et al., 2012). Curvature values range between −1 and 1 and are calculated from the cumulative response record by deviation from a straight line; 0 indicates a constant response rate throughout the interval. Curvature indices are resistant to differences in response rate, smoothing, or binning. Second, we modeled each response using generalized linear mixed-effects models (GLMM; *fitglme.m* in MATLAB) where the outcome is response time, the predictor variable was Day, and the random effect was animal. For two-interval trials, single-trial analyses were used to find start times and coefficients of variation for FI3 and FI12 trials (Church et al., 1994). All data and statistical approaches were reviewed by the Biostatistics, Epidemiology, and Research Design Core (BERD) at the Institute for Clinical and Translational Sciences (ICTS) at the University of Iowa.

Analyses of neuronal activity and basic firing properties were carried out using NeuroExplorer (Nex Technologies, Littleton, MA) and custom routines for MATLAB, as described in detail previously (Emmons et al., 2019, 2017; Parker et al., 2014). Peri-event rasters and time-histograms were constructed around houselight and lever press. As in our past work, neuronal modulations were quantified in two ways. First, we used principal component analysis (PCA), a data-driven set of orthogonal basis vectors that captures patterns of activity in multivariate neuronal ensembles. PCA was calculated from average peri-event time histograms computed from kernel-density estimates (*ksdensity.m*; bandwidth 0.5) and normalized using zscore. As in our past work, we used absolute values of PC1 scores (indicated by |PC1|) compare ramping strength across areas and days (Emmons et al., 2017; Kim et al., 2017; Parker et al., 2017, 2014).

We also used *fitglme.m* to construct GLMMs to analyze neuronal modulations. For neuron-by-neuron analysis, we used GLMMs where the outcome variable was the firing rate (binned at 0.1 s) and the predictor was time in the interval (0 to 3 seconds on FI3, or 0 to 12 seconds on FI12), interval (FI3 or FI12), or responses. Neurons with a main effect of time were considered ‘ramping’ neurons, neurons with a main effect of interval-type were considered interval-modulated, and neurons with a main effect of response were considered response-modulated. Neurons with time-related ramping were defined as those with a main-effect of time in the interval via GLMMs where the outcome was firing rate binned at 0.1 s, and the predictor was time in the interval. We also performed trial-by-trial GLMMs for all trials and neurons where the outcome was firing rate, and the predictors were area (MFC or DMS) or Day 0 or Day 1, and random effects were lever presses and neurons (Table 1). To examine interval-related modulation for all trials and neurons, we used an equation where the outcome was firing rate, the predictors were interval type (FI3 or FI12), area, or Day, and random effects were lever presses and neurons (Table 2). For two-interval performance, we used GLMMs for FI3 trials (Table 3) and FI12 trials (Table 4) on Days 1-3. Poisson distributions were used for all firing rate models.

**Table 1:** Day 0-Day 1 effects on firing rate in MFC and DMS. Model: FiringRate∼Times*Area*Learning+(1|Response)+(1|Neurons) Obs 2294280 Model R^2^ 0.19

| *Predictor* | *F* | *p* |
| --- | --- | --- |
| Times | 111.69 | **4.00E-26** |
| Area | 6.06 | **0.014** |
| Learning | 1.77 | 0.183 |
| Times:Area | 686.31 | **3.00E-151** |
| Times:Learning | 96.96 | **7.00E-23** |
| Area:Learning | 1.39 | 0.238 |
| Times:Area:Learning | 97.5 | **5.00E-23** |

**Table 2:** Firing rate on FI3 vs. FI12 trials. Model: FiringRate∼Area*Learning*Interval+(1|Response)+(1|Neurons) Obs 3018780 Model R^2^ 0.15

| *Predictor* | *F* | *p* |
| --- | --- | --- |
| Area | 0.48 | 0.49 |
| Learning | 0.58 | 0.45 |
| Interval | 3.89 | **0.05** |
| Area:Learning | 0.81 | 0.37 |
| Area:Interval | 0.32 | 0.57 |
| Learning:Interval | 13.16 | **0.0003** |
| Area:Learning:Interval | 6.35 | **0.01** |

**Table 3:** FI3 Day 1-Day 3 effects on firing rate. Model: FiringRate∼Times*Area*Learning+(1|Response)+(1|Neurons) Obs 533580 R^2^ 0.15

| *Predictor* | *F* | *p* |
| --- | --- | --- |
| Times | 0 | 0.959 |
| Area | 0.5 | 0.481 |
| Learning | 0.56 | 0.456 |
| Times:Area | 5.41 | **0.02** |
| Times:Learning | 0.04 | 0.846 |
| Area:Learning | 0.69 | 0.407 |
| Times:Area:Learning | 1.2 | 0.274 |

**Table 4:**
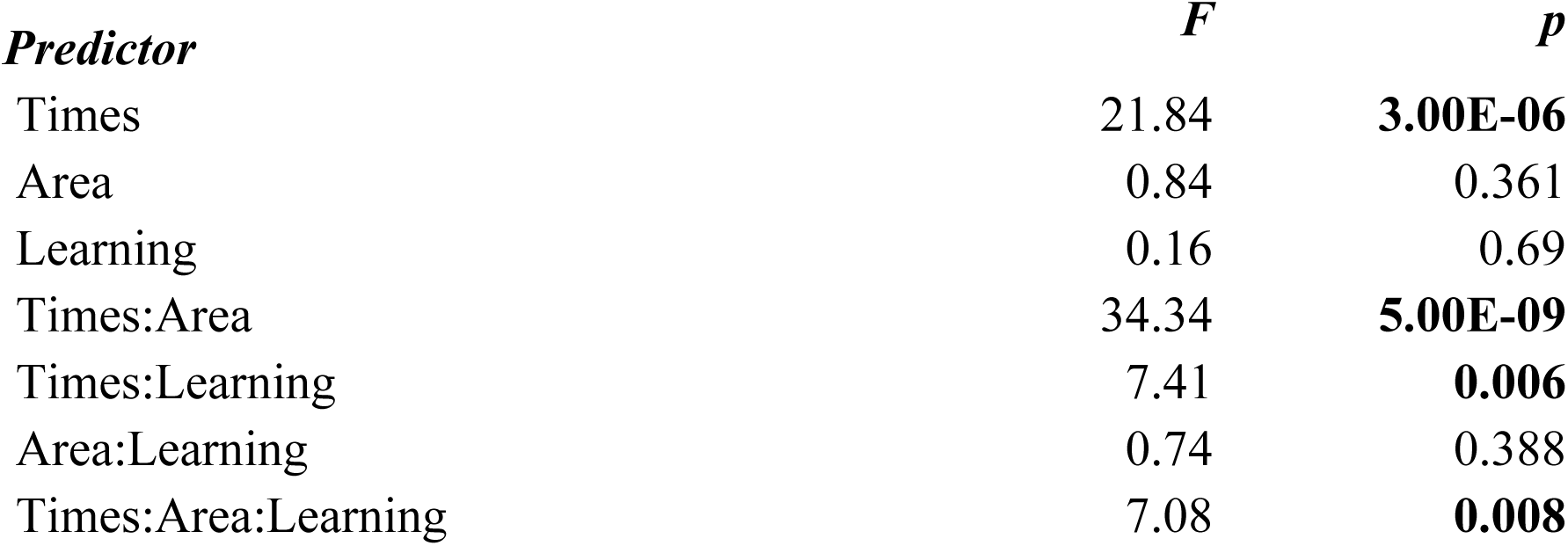
FI12 Day 1-Day 3 effects on firing rate. Model: FiringRate∼Times*Area*Learning+(1|Response)+(1|Neurons) Obs 2485200 R^2^ 0.16

| *Predictor* | *F* | *p* |
| --- | --- | --- |
| Times | 21.84 | **3.00E-06** |
| Area | 0.84 | 0.361 |
| Learning | 0.16 | 0.69 |
| Times:Area | 34.34 | **5.00E-09** |
| Times:Learning | 7.41 | **0.006** |
| Area:Learning | 0.74 | 0.388 |
| Times:Area:Learning | 7.08 | **0.008** |

We used a naïve Bayesian classifier to examine neuronal ensemble decoding, as we have in our past work (Emmons et al., 2017; Kim et al., 2017). We calculated kernel density estimates (bandwidth: 1.2) of trial-by-trial firing rates from MFC and DMS neurons. To prevent edge effects that might bias classifier performance, we included data from 6 seconds prior to trial start and 6 seconds after interval end. We used leave-one-out cross-validation to predict objective time from firing rate within a trial. We evaluated classifier performance by computing the R^2^ of objective time and predicted time only for bins during the interval. With perfect classification, the R^2^ would approach 1. Classifier performance was compared to ensembles with time-shuffled firing rates. For each area and interval, performance was compared via GLMMs of R^2^ vs. each day.

## Results

We studied temporal learning in the MFC and DMS by introducing a new 3-s fixed interval (FI3) to rats after they had been trained on a task with 12-second fixed-intervals (FI12; Fig. 1A). We compared behavior on FI12 trials only in sessions with one interval (“Day 0”) to two-interval sessions in which FI12 trials were randomly intermixed with FI3 trials (“Day 1”). We quantified the temporal control of action via a ‘curvature’ index calculated from the cumulative distribution of time-response histograms. We have used this index extensively in the past (Emmons et al., 2019; Narayanan et al., 2012). On FI12 trials, the curvature index trended towards being lower on Day 1 vs. Day 0 (Fig. 2A-B; Day 0 curvature: 0.29 ± 0.04, mean ± SEM; Day 1 curvature: 0.17 ± 0.06; signrank *p* = 0.08; Cohen’s d = 0.86). Response times were significantly shorter on Day 1 vs. Day 0 (Fig. 2C-D; Day 0: 10.14 ± 0.09 vs. Day 1: 9.61 ± 0.10 seconds; main effect of Day: F_(1,3784)_ = 12.28, *p* = 0.0005; R^2^ = 0.003). These results suggest that the timing of FI12 responses was shifted earlier on FI12 trials on Day 1 when they were intermixed with FI3 trials compared to Day 0, when FI12 trials were presented alone.

**Figure 2:**
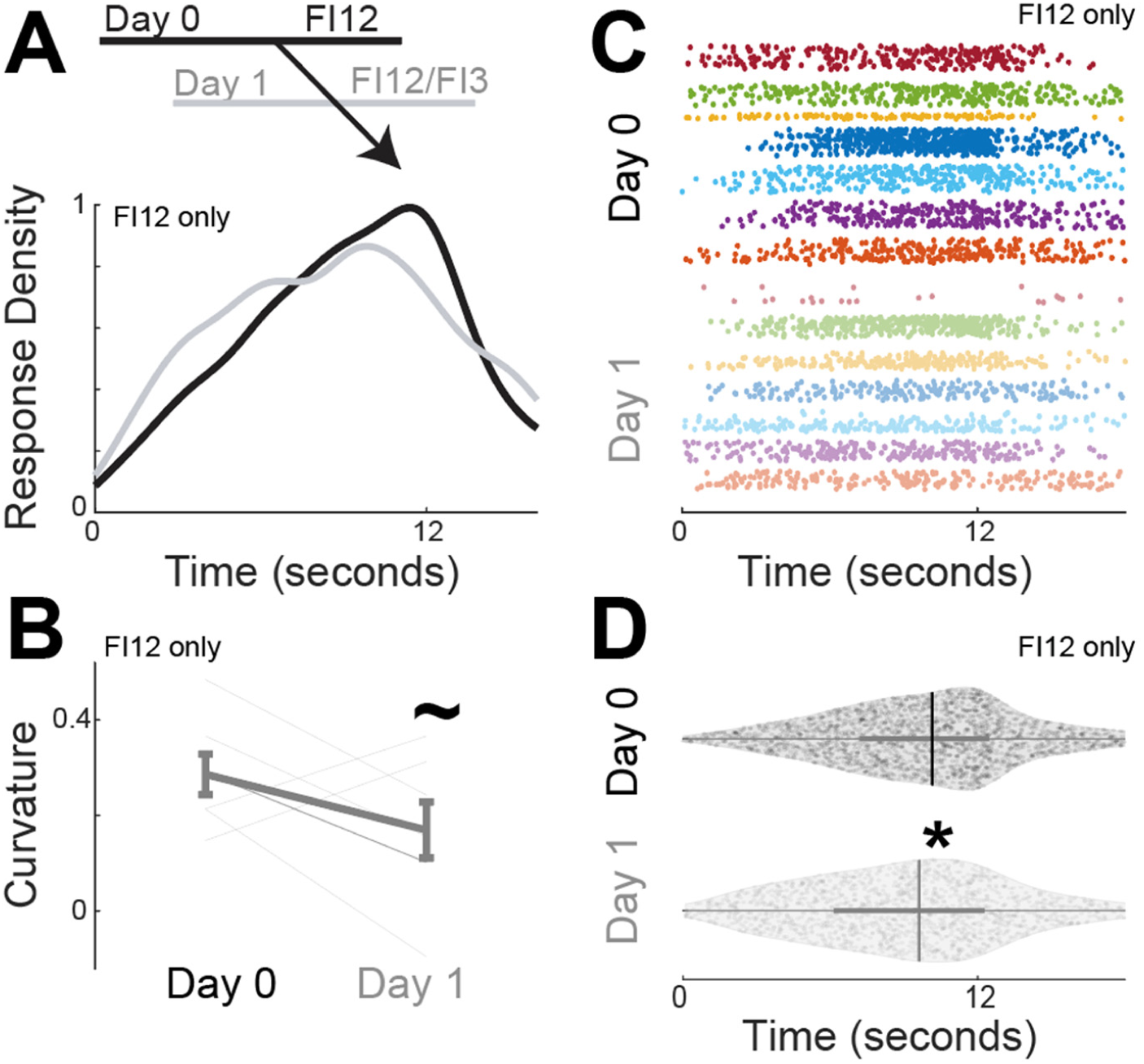
Response times reflect temporal context. A) Kernel density estimates of time-response histograms across animals on FI12 trials on Day 0 (black) when FI12 trials were presented alone to Day 1, with FI12 trials were presented alongside randomly intermixed FI3 trials. B) Curvature indices of time-response histograms from 7 animals on Day 0 vs. Day 1. C) Compilation of every FI12 response from every animal on Day 0 (darker colors) vs. Day 1 (lighter colors); each animal is represented by dots of a different color. D) Violin plot of all responses; vertical lines denote the mean and thicker horizontal gray lines span the interquartile range. Data from FI12 trials from seven animals; ∼ indicates a trend via Wilcoxon rank-sum; * indicates *p* < 0.05 via GLMMs.

We recorded neuronal ensembles simultaneously in the MFC and DMS as animals well-trained on FI12 trials on Day 0 learned to perform a two-interval task with FI3 and FI12 trials randomly intermixed on Day 1. As in our past work, we found that neurons in both brain regions exhibited time-dependent ramping, i.e. monotonic increases in firing across the interval (Emmons et al., 2017; Fig. 3A-B). On FI12 trials 35 of 59 (59%) MFC neurons exhibited ramping activity on Day 0, surprisingly, there were only 19 of 47 (40%) ramping neurons in MFC on Day 1 (Fig. 3C; X^2^ = 3.74, *p* = 0.05). In the DMS, 32 of 67 (48%) neurons ramped Day 0 and 29 of 58 (50%) ramped on Day 1, (X^2^ = 0.06, *p* = 0.80).

**Figure 3:**
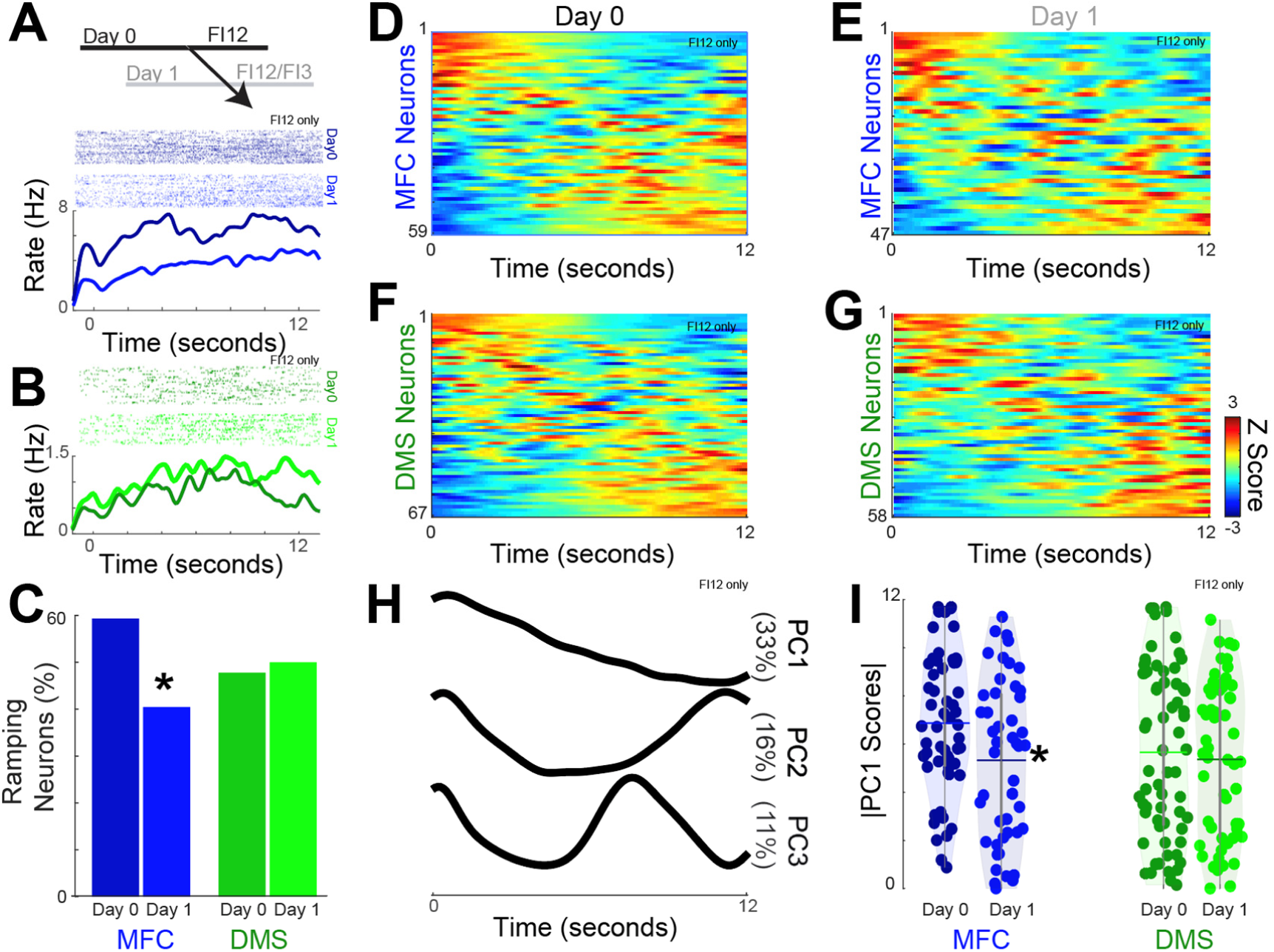
MFC ramping reflects temporal context. Peri-event rasters for single neurons in the A) MFC (blue) and B) DMS (green). Top panels: each row represents a trial; each tick is an action potential; darker colors represent Day 0 (FI12) and lighter colors represent Day 1 (FI12/FI3); all data are from FI12 trials only. C) Quantification of neurons that underwent time-related ramping via GLMMs in the MFC and DMS for Day 0 and Day 1; * indicates *p* < 0.05 via a chi-squared test. Peri-event histograms from all neurons in the MFC on Day 0 (D) and Day 1 (E) and all MSNs in the DMS on Day 0 (F) and Day 1(G). H) Principal component analyses revealed three main components; percentage of variance is indicated in parentheses. I) |PC1 scores| for MFC and DMS ensembles on Day 1 vs. Day 0. Each circle represents the |PC1 score| from a single neuron; horizontal lines denote the mean and thicker vertical lines span the interquartile range. * indicates *p* < 0.05 via Wilcoxon sign-rank. Data from MFC and DMS recordings in 7 animals.

To further compare ramping activity, we turned to principal component analysis (PCA) as a data-driven approach to compare neuronal activity patterns (Chapin and Nicolelis, 1999; Emmons et al., 2017; Narayanan and Laubach, 2009). Consistent with our prior work, we found that principal component 1 (PC1) exhibited time-related ramping (Fig 3H; Emmons et al., 2017; Kim et al., 2017; Parker et al., 2015, 2014; Zhang et al., 2019). For the MFC, the strength of |PC1| was lower on Day 1 than Day 0 (Fig. 3I; signrank *p* = 0.03; Cohen’s d = 0.50), whereas there was no consistent difference for the DMS (Fig 3I; signrank *p* = 0.60). Trial by-trial analysis of firing rate on FI12 trials revealed that time-related ramping interacted with both the brain area (MFC and DMS) as well one- vs. two-interval sessions (i.e., Day 0 vs. Day 1). There was a three-way interaction between time-related ramping, brain area, and one vs. two intervals (Table 1). For FI12 trials, these data suggest that time-related ramping in the MFC was stronger on Day 0 compared to Day 1 when FI12 trials were intermixed with FI3 trials. Our results suggest that neuronal ensembles in the MFC, but not in the DMS, are sensitive to the temporal context (Jazayeri and Shadlen, 2010; Shi et al., 2013).

Next, we compared MFC and DMS activity on FI3 and FI12 trials. First, we analyzed fixed-interval behavior using single-trial analysis, which was developed for peak-interval timing but can be useful to analyze start times during fixed-interval tasks (Church et al., 1994; Emmons et al., 2019). On the first day of the shorter 3-second interval, single-trial analysis revealed that animals had shorter start times on FI3 trials compared to FI12 trials (FI3: 3.18 ± 0.69 vs. FI12 7.06 ± 0.34, signrank *p* = 0.03; Cohen’s d = 2.9; single-trial analysis could not compute FI3 start times from one animal, which was subsequently removed from this analysis). One indication that timing processes are scalar is that the coefficient of variation (CV—the ratio of standard deviation of temporal estimates to the mean) is relatively constant at different intervals (Gibbon et al., 1984; Rakitin et al., 1998). Accordingly, we found that during fixed-interval performance, start time CVs were similar on FI3 and FI12 trials (FI3: 0.59+/-0.12 vs. FI12: 0.41+/-0.03 signrank p: 0.56). These data suggest that start times during fixed-interval timing exhibit scalar properties (Gibbon et al., 1984).

Ramping neurons can have distinct slopes of firing rates vs. time on FI3 and FI12 trials (Fig 4A). We ran GLMMs where firing-rate slope vs. time was the outcome variable, and FI3 vs. FI12 interval and Day were predictor variables; note that we were interested in the magnitude of the slope and we focus on its absolute value, indicated by |slope|. Consistent with past work by our group and others, ramping neuron |slopes| were consistently steeper on FI3 vs. FI12 trials for both the MFC (Fig. 4B; main effect of interval: F_(143)_ = 4.52, *p* = 0.04, R^2^ = 0.22) and for the DMS (main effect of interval: F_(214)_ = 6.91, *p* = 0.01; R^2^ = 0.40(Emmons et al., 2017; Mello et al., 2015; Wang et al., 2018). There was no effect of Day or higher interactions for either MFC or DMS. These data are consistent with influential drift-diffusion models of interval timing, suggesting that drift rates increase with shorter intervals (Simen et al., 2011).

**Figure 4:**
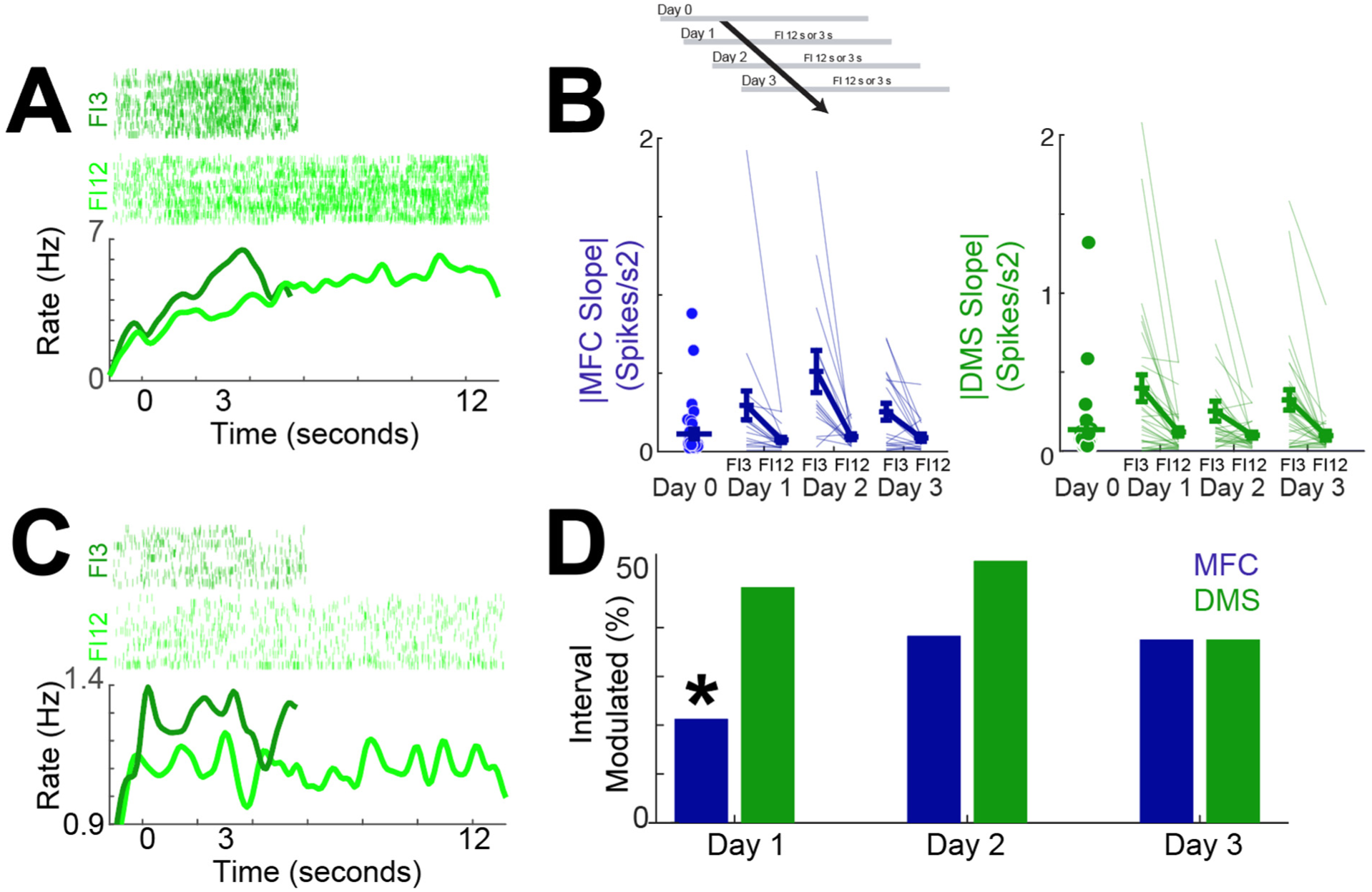
MFC and DMS activity is distinct on FI3 vs. FI12 trials. A) An exemplar ramping neuron from the DMS; the slope of firing rate vs. time was steeper on FI3 vs. FI12 trials. B) |Slopes| for MFC (blue) and DMS (green) neurons on Day1, Day 2, and Day 3 for FI3 and FI12 trials; for both MFC and DMS |slopes| were higher on FI3 trials. C) An exemplar neuron from the DMS that fired differentially on FI3 vs. FI12 trials. D) The number of neurons with a main effect of firing rate vs. interval for the MFC and DMS. * indicates *p* < 0.05 via a chi-squared test; Data from MFC and DMS recordings in 7 animals.

Next, we searched for neurons in which firing rates were a function of interval duration (Fig 4C). Specifically, we used GLMMs to identify neurons with a main effect of interval on firing rate (Fig. 4E; FI3 vs. FI12 trials). Interval-modulated neurons were more common in the DMS than the MFC on Day 1 of two-interval performance (Fig 4D; MFC 10 of 47 vs. DMS: 28 of 58; X^2^ = 8.20; *p* = 0.004). Interestingly, ∼50% of interval-modulated neurons also had ramping activity in MFC (5 of 10) and DMS (15 of 28). Of note, the number of interval-modulated neurons was not different between MFC and DMS on Days 2 and 3 (Fig 4D). Consistent with these analyses, GLMMs revealed a significant interaction between interval-modulation, brain area, and Days 1-3 (Table 2).

A comparison of behavior across the three days of two-interval performance revealed that response times shortened for both FI3 and FI12 trials (Fig. 5A-B: GLMM FI3: F_(1741)_ = 8.35, *p* = 0.0002; R^2^ = 0.05; FI12: F_(4796)_ = 6.98, *p* = 0.0009; R^2^ = 0.002), but the curvature of time-response histograms did not reliably change (FI3: F_(40)_ = 0.03, *p* = 0.87, R^2^ = 0.17; FI12: F_(40)_ = 1.05, *p* = 0.31, R^2^ = 0.61). As in prior work demonstrating temporal scaling in the MFC and DMS, PCA revealed that the principal components for FI3 and FI12 trials were very similar (Fig. 5C-D; Pearson’s *rho* correlation: PC1: = 0.99 *p* < 0.001; PC2: −0.95, *p* < 0.001; PC3: 0.95, *p* < 0.001; (Emmons et al., 2017; Mello et al., 2015; Wang et al., 2018). However, |PC1| did not change consistently over the three days of two-interval performance for MFC or DMS (Fig. 5E-H; Table 3-4). Taken together, these data indicate that ramping-related patterns of activity of corticostriatal ensembles did not consistently change as animals performed two-interval tasks on Days 1-3.

**Figure 5:**
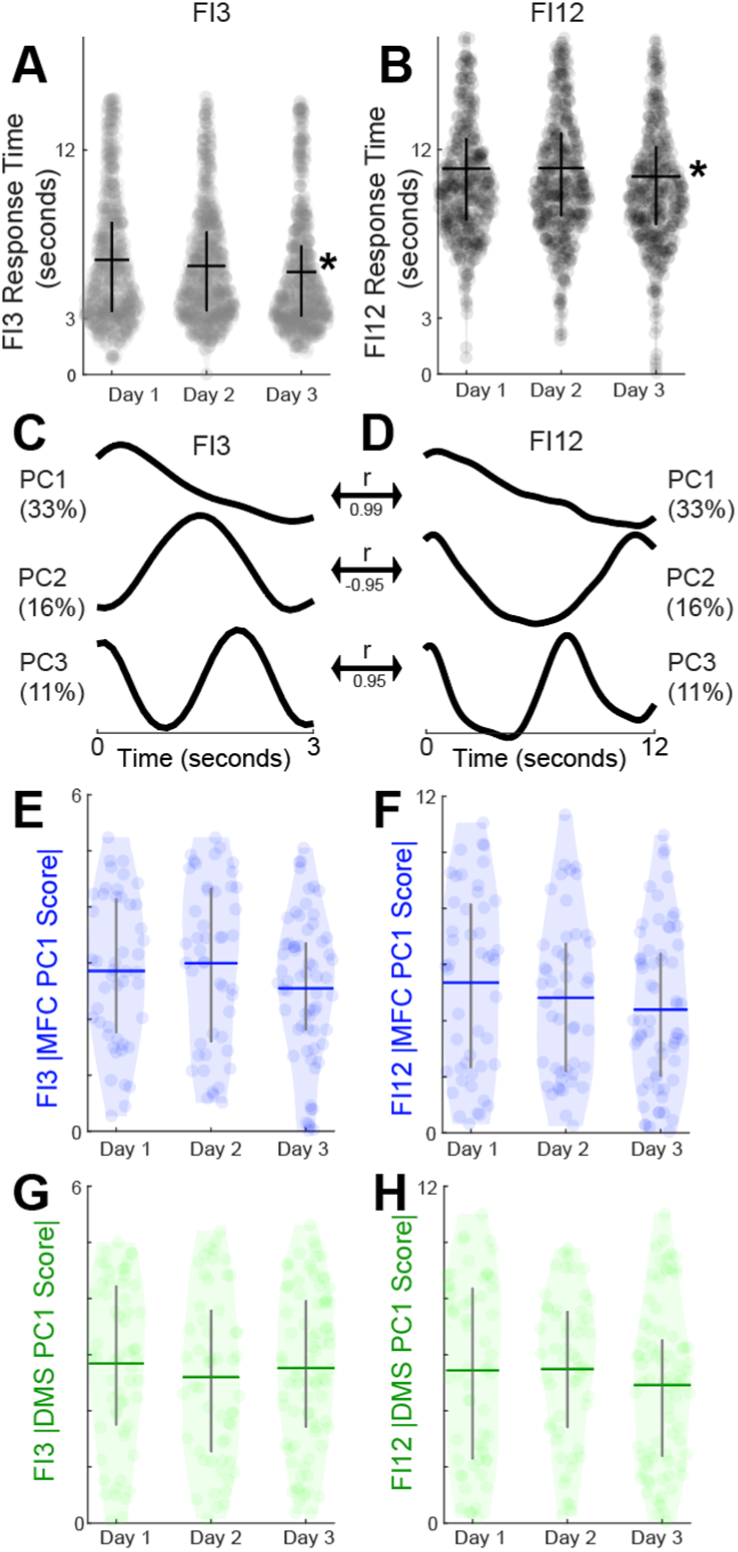
MFC and DMS ramping is stable with two-interval performance. A) Response times by day for FI3 and B) FI12 trials over the three days of two-interval performance. Principal components for C) FI3 and D) FI12 trials. Ramping activity as measured by |PC1| scores for E) FI3 trials in the MFC (blue), F) FI12 trials in the MFC, G) FI3 trials in the DMS, and H) FI12 in the DMS. Horizontal lines denote the mean and thicker vertical gray lines span the interquartile range. * indicates *p* < 0.05 via GLMMs. Data were from MFC and DMS recordings in 7 animals.

We turned to decoding analyses based on machine learning to capture more complex features of MFC and DMS ensembles (Fig 6; Emmons et al., 2017; Gouvea et al., 2015; Kim et al., 2017). Specifically, we constructed neuron-dimensional arrays of smoothed trial-by-trial firing rates over the interval binned at 0.1 seconds. We decoded time in the interval from ensemble firing rates using naïve Bayesian classifiers. Classifier performance was assessed by computing the variance explained (R^2^) of predicted vs. observed time. For all sessions, R^2^ was much less for time-shuffled ensembles—i.e., ensembles constructed from neuronal activity shuffled in time (Fig. 6B-G; signrank *p* = 4*10^−47^; Cohen’s d = 1.57). We found that temporal decoding had a main effect of Day only for the DMS Ensembles on FI3 trials (Fig. 6F&G; F_(108)_ = 6.07, *p* = 0.02, R^2^ = 0.06). These results suggest that as the response times shortened with two-interval performance, temporal decoding improved only for DMS ensembles on FI3 trials.

**Figure 6:**
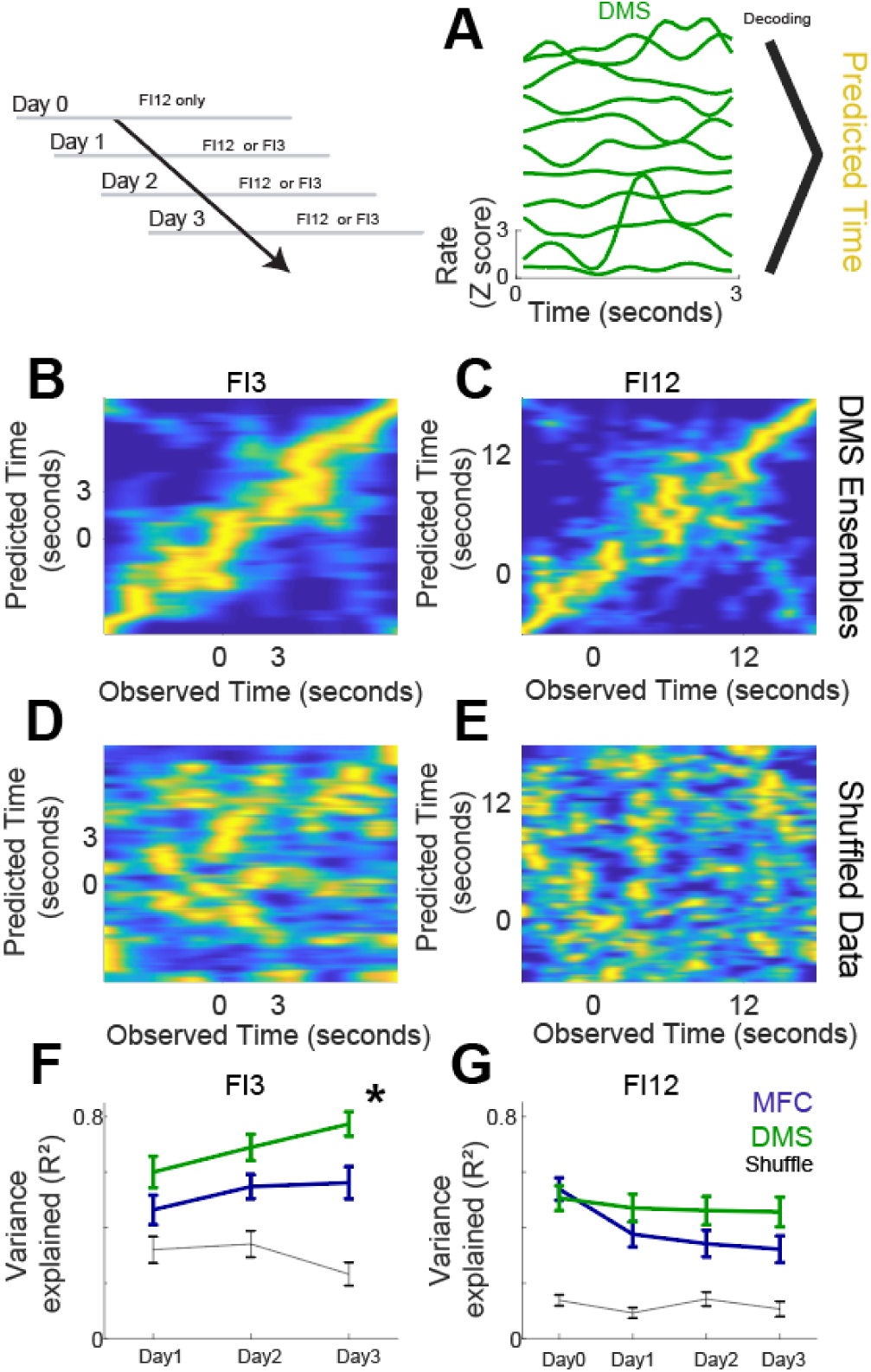
Decoding reveals that DMS improves temporal predictions with two-interval performance. A) We trained decoders (naïve Bayesian classifiers) to predict time from firing rate. B) Decoder performance for DMS ensembles for FI3 trials and C) for FI12 trials. Predicted time is on the y axis and observed time is on the x axis; decoded time is in yellow. D) Decoded performance for the same DMS ensembles with time-shuffled data for FI3 trials and E) FI12 trials. F). We measured decoder performance by calculating the variance explained (R^2^) of predicted vs. observed time. For FI3 trials, DMS ensembles increased R^2^ with two-interval performance, while MFC and G) FI12 MFC and DMS decoding was unchanged. * indicates a main effect of Day 1 → Day 3 via GLMMs; data from MFC and DMS recordings in 7 animals.

## Discussion

We found three key distinctions between MFC and DMS during temporal learning. First, time-related ramping in the MFC decreased as animals that had been trained on a one-interval task learned to respond to a second interval that was novel and shorter. Second, interval-modulated neurons were more common in the DMS early in two-interval performance. Third, time-related ramping in the MFC and DMS did not change with two-interval performance, but temporal decoding improved only for the DMS on FI3 trials. Our data suggest that MFC ensembles are sensitive to the context or ‘rules’ in the task – i.e., FI12 vs. FI3/FI12, while the DMS optimizes behavior particularly on FI3 trials. These data provide insight into the relative roles of prefrontal and striatal networks during temporal learning.

These results contradicted our hypothesis that time-related ramping in the MFC and DMS would be similarly affected by the introduction of a new temporal interval. Our hypothesis was based on five lines of evidence: 1) the existence of strong projections from the MFC to the DMS (Gabbott et al., 2005; Han et al., 2017; Wall et al., 2013), 2) clear roles for both structures in interval timing (Coull et al., 2011; De Corte et al., 2019; Emmons et al., 2017; Meck, 2006), 3) similarities in time-related ramping in the MFC and DMS (Emmons et al., 2017), 4) the necessity of MFC activity for DMS ramping (Emmons et al., 2019, 2017), and 5) our recent demonstration that the stimulation of axons that project from the MFC to the DMS is sufficient to increase time-related ramping in the DMS (Emmons et al., 2019). Given these data, it is notable that the MFC and DMS play distinct roles during temporal learning, although this observation is concordant with the vastly different connectivity and synaptic organization of these two structures (Shepherd, 2003). Nevertheless, decreases in MFC ramping after the introduction of a shorter interval suggest that MFC ramping is sensitive to the temporal context of the one-interval vs. two-interval task, and they may reflect Bayesian priors of temporal probabilities (Jazayeri and Shadlen, 2010; Shi et al., 2013).

Differences between the MFC and DMS were anticipated based on a recent comparison of neuronal ensembles during a temporal categorization task that involved maze running (Kim et al., 2018). This study indicated that ramping was more prevalent in the MFC than the dorsal striatum. In this task as the intervals became longer, temporal decoding by the MFC was less effective than that by the striatum. Although this task was more complex than ours and a number of others (Bakhurin et al., 2017, 2016; Donnelly et al., 2015; Narayanan, 2016; Wang et al., 2018), the strong temporal encoding across corticostriatal ensembles is consistent with our findings here.

We found that the DMS contained more neurons in which there was a main effect of interval compared to the MFC on the early days of two-interval performance. Notably, half of interval-modulated neurons were not ramping. These data suggest that patterns beyond time-related ramping encode information about temporal intervals. On progressive days of two-interval performance, interval-related activity between the MFC and DMS equalized. Because our task design involved a second cue for FI3 intervals, we cannot distinguish whether this activity was related to working memory for temporal intervals, cue-related processing, or other aspects of interval timing. Future work using more advanced learning paradigms may clarify these patterns of activity.

Our findings are in line with drift-diffusion models of two-interval tasks, as we find that time-related ramping scales with the interval duration (Simen et al., 2011). We find that MFC ramping is sensitive to temporal context whereas DMS ramping is not, and that non-ramping interval-related modulations and temporal predictions in the DMS change with two-interval performance. These results suggest that time-related ramping reflects distinct processes in MFC and DMS. Given that MFC activity influences ramping in DMS (Emmons et al., 2019, 2017), DMS ramping activity might integrate aspects of MFC ramping as well as non-ramping activity.

Because time-related ramping activity in MFC and DMS ensembles did not change during two-interval performance, ramping activity may be remarkably stable in both brain regions when the temporal context does not change. It is unclear how ramping might change with extended periods of behavior over several days or weeks (Barnes et al., 2005; Graybiel, 2008; Yin et al., 2005). However, we did find that on FI3 trials, temporal decoding in the DMS improved even though DMS ramping was stable. In the DMS patterns beyond ramping activity might change during two-interval performance and contribute to improved temporal decoding (Paton and Buonomano, 2018). The improvement in temporal prediction despite unchanged ramping activity supports improved ‘population clock’-based temporal predictions during FI3 trials (Karmarkar and Buonomano, 2007; Laje and Buonomano, 2013).

To our knowledge, our study is one of the first to record from corticostriatal ensembles during temporal learning. The striatum has a well-established role in learning other contexts including habit formation, reversal learning, and instrumental learning (Graybiel and Grafton, 2015; Kimchi and Laubach, 2009; Yin and Knowlton, 2006). However, direct comparisons of cortical and striatal learning in rodents, in any context, are rare. One exception is a recent study of prenatal alcohol exposure, which showed that the orbitofrontal cortex disengages and the dorsal striatum updates reward contingencies (Marquardt et al., 2020). These findings parallel the changes in the MFC and DMS that we report here. The observation that prenatal exposure to alcohol leads to changes in cortical activity underscores the clinical significance of this brain circuit.

During associative learning in primates, corticostriatal ensembles are highly sensitive to learning, with striatal neurons rapidly encoding new associations and the prefrontal cortex learning more slowly (Pasupathy and Miller, 2005). Primate striatal neurons rapidly encoded stimulus-response associations, whereas primate prefrontal neurons encoded category abstraction (Antzoulatos and Miller, 2011). In line with these results, we found that time-related ramping decreased in two-interval vs. one-interval sessions, suggesting that prefrontal ensembles may be sensitive to temporal categories or context. It is important to note that these primate studies recorded from lateral prefrontal areas, which lack a clear rodent analogue (Laubach et al., 2018), and that they employed vastly different task conditions. Nevertheless, our work provides insight into the dynamics of rodent corticostriatal ensembles during an elementary temporal learning paradigm.

Our study has several limitations. First, we used fixed-interval timing; peak-interval timing tasks might enable more precise dissection of start and stop times (Rakitin et al., 1998). Second, our techniques cannot identify the genetic or molecular identity of recorded neurons. This detail would be of particular interest in the case of the DMS, which contains D1 and D2 MSNs (Kreitzer, 2009). Third, we are unsure if the MFC and DMS neurons we captured were connected, because of the sparsity of cortical projections and constraints of our recording techniques (Wall et al., 2013). This limitation might be overcome in future work by exploiting optogenetic tagging and retrograde viral tracing to isolate corticostriatal projections (Otis et al., 2017). Studying how MFC-DMS connectivity changes with learning might provide further insight in corticostriatal circuits. Fourth, we cannot reliably follow neurons over separate sessions. Finally, we were unable to clearly identify clear correlates of temporal learning during the first two-interval session on Day 1. Corticostriatal ensembles may rapidly learn the new interval. Such rapid learning has been observed in the striatum, but capturing it might require a different task design to capture trial-by-trial neuronal dynamics during learning (Kimchi and Laubach, 2009).

In summary, we investigated corticostriatal ensembles in rodents that had been well-trained to perform a single fixed-interval timing task while they learned to incorporate a new interval. We discovered that time-related ramping activity in the MFC decreased following introduction of the shorter interval, whereas ramping activity in the DMS was unchanged. We also found that more DMS neurons fired differentially on each interval compared to the MFC early in two-interval performance. Finally, corticostriatal ramping activity did not change on the days following the initiation of two-interval performance, yet DMS temporal decoding improved. Taken together, our data suggest that the MFC and DMS play distinct roles in temporal learning.

## Acknowledgements

We thank Morgan Kennedy and Tomas Lence for technical help.

## Contributions

EE, YK, and NN designed experiments; EE and GTC collected data, EE, YK, KC, and NN analyzed data, AB independently checked the code and data, and EE, GTC, MW, AB, YK, and NN wrote the manuscript.

## Data and Code

Available at narayanan.lab.uiowa.edu

